## Supplementary figures, tables and methods for "An ancestral apical brain region contributes to the central complex under the control of *foxQ2* in the beetle *Tribolium castaneum*"

#### 1. Supplementary Figures

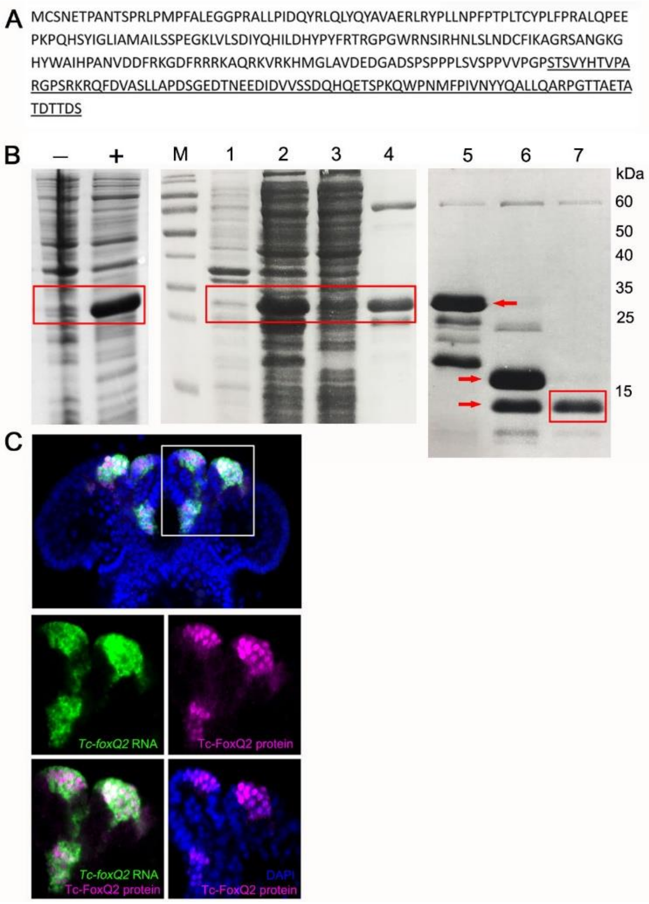


**Fig. S1. Generation of a Tc-FoxQ2 antibody**. (A) Protein sequence of Tc-FoxQ2. The C-terminus containing 85 amino acids (underlined) has little homology to other proteins in *Tribolium* and was used for protein expression. (B) Coomassie-blue stained SDS-PAGE gel analysis of expression and purification of Tc-FoxQ2. (-) Before IPTG induction; (+) after IPTG induction. M, marker; lane 1, cell pellet; lane 2, supernatant; lane 3, flow through after $Ni^{2+}$ chelate affinity chromatography; lane 4, eluted fractions by imidazole; lane 5, before SUMO protease digestion (red arrow); lane 6, after SUMO protease digestion, two bands are observed (red arrows): 6xHis-SUMO and Tc-FoxQ2; lane 7, flow through after re-$Ni^{2+}$ chelate affinity chromatography which contains Tc-FoxQ2. (C) Expression of *Tc-foxQ2* RNA (green) and Tc-FoxQ2 protein (magenta) in the embryo. *Tc-foxQ2* RNA is detected throughout the cytoplasm, while Tc-FoxQ2 protein is detected in the nuclei (blue). *Tc-foxQ2* RNA and Tc-FoxQ2 protein show a high overlap.

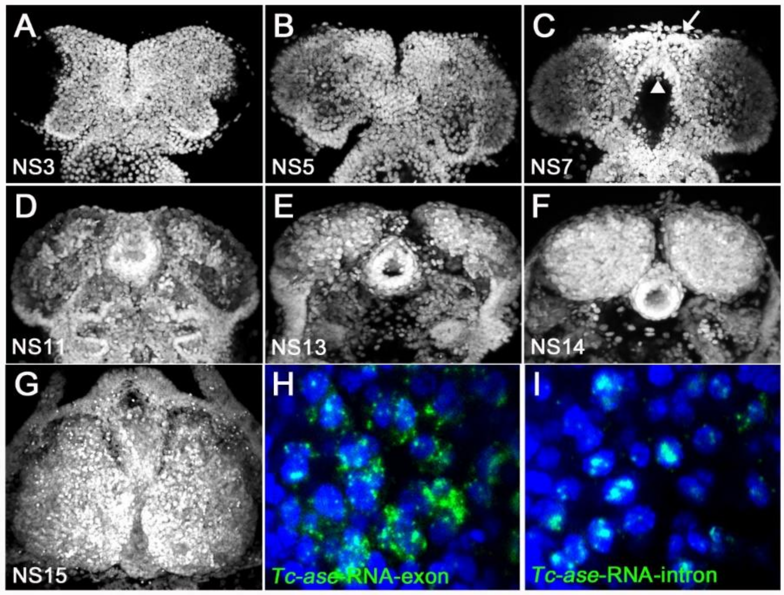


**Fig. S2. Developmental staging and comparison of exonic versus intronic *Tc-asense* probes.** In *Tribolium*, the established morphological staging system (Biffar and Stollewerk, 2014) with respect to early neurogenesis has 15 stages, termed NS1 to NS15. (A, B) From NS3 to NS5, the neuroectoderm is still a flat sheet of cells. (C) At NS7, the labral and stomodeal buds have formed (arrow, white arrowhead). (D) At NS11, the brain grows thicker and contains an increased number of cells. (E, F) From NS13 onwards, the brain hemispheres become an oval shape and grow. (G) At NS15, the brain hemisphere shows a pear-like shape. All planes are dorsal view. (H) *Tc-ase*-RNA exonic probe marks cytoplasm. (I) *Tc-ase*-RNA intronic probe marks only nuclei, which allows cell identification.

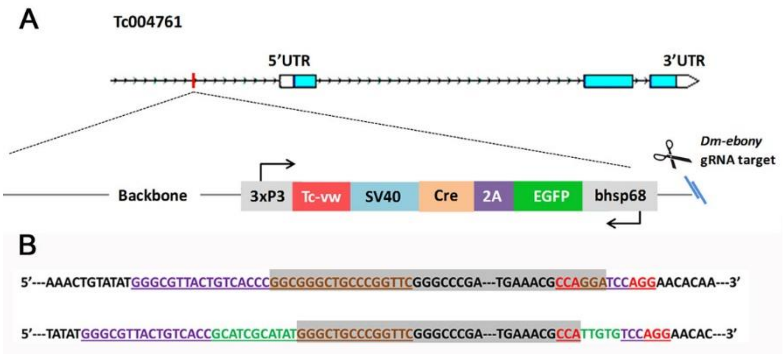


**Fig. S3. Characterization of the *foxQ2-5'-line*.** (A) The insertion (including the plasmid backbone) is about 6.8 kb in length and the bhsp68 promoter is located approximately ~700 bp upstream of the transcription start site. The EGFP transcription unit is oriented in the opposite direction to *Tc-foxQ2*. (B) The upper row is the predicted sequence of a perfect insertion without indels where grey indicates the left and right extremes of the 6.8kb insert and the gRNA target sequence is underlined. The lower row is the actual sequencing results of this line. The insertion occurred at the predicted site (3 bp upstream of the PAM, AGG shown in red) and small insertions occurred at both integration sites (green). The purple bases represent the gRNA1 target sequence (targeting the genomic DNA) and the brown bases represent the gRNA-eb target sequence (targeting the plasmid for linearization). The genomic region and sequence of *Tc-foxQ2* (*Tc004761*) are viewed by the iBeetle genome browser (http://ibeetle-base.uni-goettingen.de/gb2/gbrowse/tribolium/).

gRNA1 target sequence: 5' GGGCGTTACTGTCACCCTCCAGG 3'

gRNA-eb target sequence: 5' CCAGGAGGCGGGCTGCCCGGTTC 3'

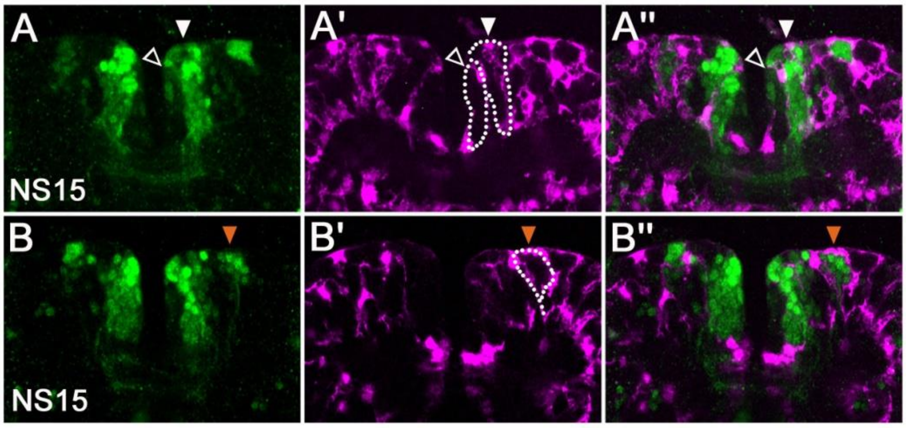


**Fig. S4. *Tc-foxQ2* cell clusters are surrounded by glial sheets.** A cross of the *foxQ2-5′-line* (green) with the *glia-line* (Koniszewski et al. 2016; magenta) was imaged. (A-A′′) Most of the cells of the *anterior-median-foxQ2-cluster* are within one glia sheet (white arrowheads, dashed line). The medial-most cells which show comparably weak EGFP signal are surround by another glia sheet (open arrowheads, dashed line). (B-B′′) The *anterior-lateral-foxQ2-lineage* is surrounded by one glia sheet (orange arrowheads, dashed line).

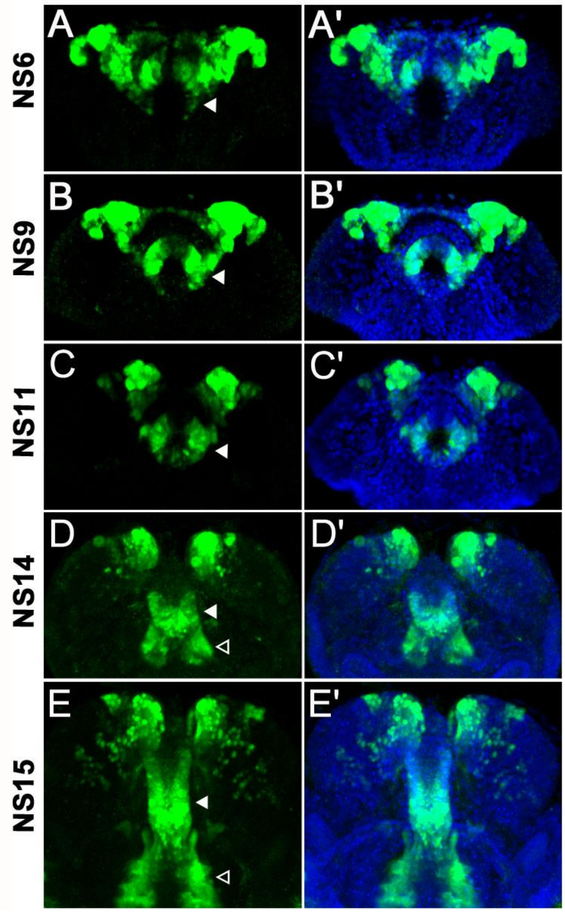


**Fig. S5. *foxQ2-5′-line* marks cells of the stomodeum and the lateral parts of the labrum.** Immunohistochemistry visualizes the EGFP (green) expressed by the *foxQ2-5′-line.* The is visualized with DAPI staining (blue). The expression of *Tc-foxQ2* marked by EGFP from NS6 to NS15 is shown. (A-C′) The stomodeum is located posteriorly in anterior median region and cells of it except for the dorsal roof are marked (white arrowheads). (D-E′) At NS14 and NS15, additional marked epidermal cells in the lateral parts of the labrum are observed (open arrowheads).

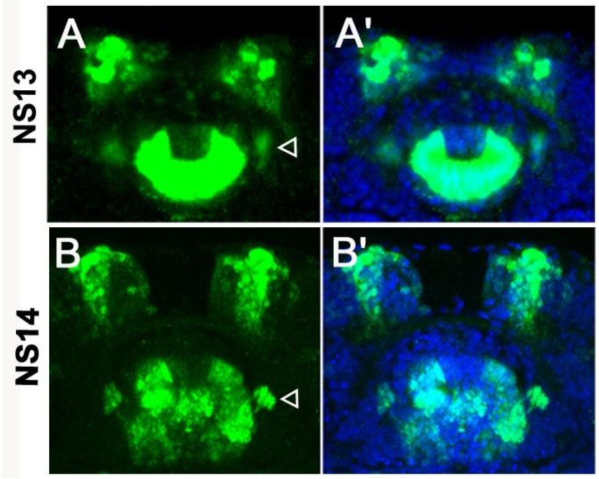


**Fig. S6. The *foxQ2-5'-line* marks cells lateral of the stomodeum.** Immunohistochemistry visualizes the EGFP (green) derived from the *foxQ2-5'-line.* The morphology of the brain is visualized with DAPI staining (blue). (A-A') At NS13, several cells lateral to the stomodeum are observed (open arrowhead). (B-B') At NS14, this group of cells project into the posterior circumesophageal commissure (open arrowhead).

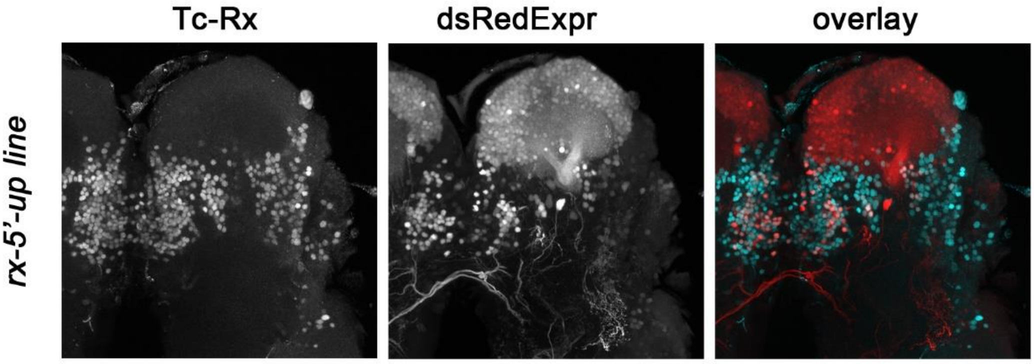


**Fig. S7. Partial co-localization of reporter protein with endogenous protein in the rx-5'-up line.**

### 2. Supplementary Tables

**Table S1. Number of Tc-FoxQ2$^{+}$ neural precursors (NPCs) per half-side during embryogenesis.**

| Stage/Individual | #1 | #2 | #3 | #4 | #5 | #6 | Mean |
|---|---|---|---|---|---|---|---|
| NS8 | 15 | 16 | 15 | 12 | 17 | 16 | 15 |
| NS11 | 10 | 9 | 9 | 11 | 12 | 10 | 10 |
| NS14 | 6 | 5 | 6 | 5 | 7 | 5 | 6 |

**Table S2. gRNAs target sequences and oligonucleotides used for generating gRNAs.** The PAM sequence is marked in red. The orange sequence represents the complementary overhangs to the vector generated by *BsaI* digestion.

| Name of gRNA | Genomic target sequence | Sense oligo | Antisense oligo |
|---|---|---|---|
| gRNA1 | GGGCGTTACTGTCACCCTCCAGG | TTCGGGCGTTACTGTCACCCTCC | AAACGGAGGGTGACAGTAACGCC |
| gRNA2 | GTGGCGGGGCGGAGCCAACGCGG | TTCGTGGCGGGGCGGAGCCAACG | AAACCGTTGGCTCCGCCCCGCCA |
| gRNA-eb | GAACCGGGCAGCCCGCCTCCTGG | TTCGAACCGGGCAGCCCGCCTCC | AAACGGAGGCGGGCTGCCCGGTT |

**Table S3. Plasmids used as PCR templates and molecular cloning.**

| Plasmid and internal number | Description |
|---|---|
| pJET1.2 | backbone for knock-in construct cloning |
| # 82 pBac[3xP3-gTc'v; Tc'ems-overex] | used to amplify 3xP3, *Tc-vw* |
| # 169 pSLfa[Tc-hsp_p-ECFP-SV40] | used to amplify SV40 |
| # 133 pSLfa[Tc'hsp5'-Cre recomb-3'UTR]fa | used to amplify Cre |
| # 65 pBac[3xP3-gTc'v; Tc'hsp-Promotor RC] | used to amplify bhsp |

**Table S4. Number of the injected embryos, the hatched larvae and developed adult beetles as well as the efficiency of germ line transmission.**

| Target region | Injected embryos | Hatched larvae | Hatching rate[%] | Developed adults | Survival rate [%] | Efficiency [%] |
|---|---|---|---|---|---|---|
| **Upstream** | 1496 | 479 | 31.99 | 234 | 15.64 | 6/234=2.6 |

**Table S5. Number of cells in the *anterior-median-foxQ2-cluster*.**

| Stage/Ind. | | #1 | #2 | #3 | #4 | #5 | Mean |
|---|---|---|---|---|---|---|---|
| **NS13** | **Total cells** | 89 | 93 | 84 | 85 | 96 | 89 |
| | **NPCs** | 12 | 11 | 13 | 14 | 13 | 12 |
| | **Weak cells** | 13 | 10 | 12 | 15 | 10 | 12 |
| **NS14** | **Total cells** | 154 | 142 | 150 | 149 | 158 | 150 |
| | **NPCs** | 11 | 15 | 14 | 13 | 16 | 14 |
| **NS15** | **Total cells** | 219 | 210 | 230 | 238 | 225 | 224 |

**Table S6. Number of cells in the *anterior-lateral-foxQ2-lineage*.**

| Stage/Ind. | #1 | #2 | #3 | #4 | Mean |
|---|---|---|---|---|---|
| **NS14** | 8 | 6 | 7 | 6 | 7 |
| **NS15** | 15 | 17 | 18 | 20 | 18 |

**Table S7. Number of cells in respective groups of *Ten-a-green* line.**

| Location/Ind. | | #1 | #2 | #3 | #4 | Mean |
|---|---|---|---|---|---|---|
| Anterior group | WT | 38 | 40 | 41 | 36 | 39 |
| | RNAi | 22 | 28 | 17 | 16 | 20 |
| Posterior-lateral group | WT | 36 | 31 | 27 | 34 | 32 |
| | RNAi | 7 | 16 | 22 | 17 | 16 |
| Posterior-median group | WT | 30 | 28 | 27 | 24 | 27 |
| | RNAi | 8 | 17 | 15 | 18 | 15 |
| Total number | WT | 104 | 99 | 95 | 94 | 98 |
| | RNAi | 37 | 61 | 54 | 51 | 51 |

**Table S8. Number of cells of *Tc-rx* line.**

| Location/Ind. | | #1 | #2 | #3 | #4 | #5 | #6 | Mean |
|---|---|---|---|---|---|---|---|---|
| Anterior-median group | WT | 35 | 42 | 43 | 36 | 41 | 44 | 40 |
| | RNAi | 11 | 9 | 7 | 11 | 12 | 6 | 9 |

**Table S9. Number of cells of *foxQ2-5'-line*.**

| Anterior-median cluster and lateral lineage | | #1 | #2 | #3 | #4 | Mean |
|---|---|---|---|---|---|---|
| NS13 | WT | 89 | 93 | 96 | 85 | 89 |
| | RNAi | 45 | 50 | 28 | 43 | 42 |
| NS15 | WT | 227 | 250 | 253 | 243 | 243 |
| | RNAi | 104 | 95 | 120 | 117 | 109 |

**Table S10. Primer sequences and purposes used for cloning.**

| Primer | Sequence ( 5'-3') | Purpose |
|---|---|---|
| BH_C_ter_fwd | CCAGGTCTCATGGTTCCACGTCCGTTTATCACAC | Cloning of *Tc-foxQ2* C-terminal fragment with *BsaI* |
| BH_C_ter_rev | GGGGGTCTCCTCGAGTTAAGAGTCTGTGGTGTCGGTGGC | |
| RT1_EGFP_bhsp_rev | TGCTCACCATGTTTGACTTTGAATTCACTAGTAAATAATTCACTCAACTTTGTTAAAG | Cloning of bhsp, EGFP, 2A and Cre |
| RT2_bhsp_EGFP_fwd | AGTGAATTCAAAGTCAAACATGGTGAGCAAGGGCG | |
| RT3_2A_EGFP_rev | GTCTCCTGCTTGCTTTAACAGAGAGAAGTTCGTGGCTCCGGATCCCTTGTACAGCTCGTCCATGCC | |
| RT4_2A_Cre_fwd | GCCACGAACTTCTCTCTGTTAAAGCAAGCAGGAGACGTGGAAGAAAACCCCGGTCCTATGTCCAATTTACTGACCGTACACCAA | |
| RT5_3xP3_fwd | GATTCTAGACATTATTCATTAGAGACTAATTCAATTAGAGCTAATTCAATTAGGATCC | Cloning of 3xP3, vermillion and Sv40 |
| RT6_Sv40_Cre_rev | AATGGAAACAATTAAGATGAGTTTGGACAAACCACA | |
| RT7_Sv40_Cre_fwd | TTGTCCAAACTCATCTTAATTGTTTCCATTCGACACGT | |
| RT8_Sv40_vw_rev | GTATGGCTGATTATGACTAATCGCCATCTTCCAGCA | |
| RT9_Sv40_vw_fwd | GGAAGATGGCGATTAGTCATAATCAGCCATACCACA | |
| RT10_ebony_fwd | GTCGGGCCCGAACCGGGCAGCCCGCCTCCTGGCGTTTCATATATAAGCGCGGTCTCG | Cloning of ebony site with *ApaI* |
| BH_AG _fwd | GCGCTGGCATTTTTAAATCACG | Testing the insertion site |
| BH_AG_rev | ATACTGTAGAGCTGGAGCC | |
| BH_EGFP_rev | TGGTGCAGATGAACTTCAG | |
| rx_fwd | ACGGATCCGGGATCAAGCGTAAATGGGACGTCCCATACAATA | Cloning of *Tc-rx* 5'up regulatory region with *BamHI*, *NheI* |
| rx_rev | ATCTACGCTAGCCTTCACAACGGTCCGATTCTATCGC | |
| DsRed_fwd | ATGGCCTCCTCCGAGGA | Cloning of DsRedExpress |
| DsRed_rev | CTACAGGAACAGGTGGTGG | |

### 3. Supplementary Material and Methods

#### Generation of a Tc-FoxQ2 polyclonal antibody

The C-terminus (amino acids 202–286) was amplified from cDNA by PCR using primer pairs with *BsaI* restriction site forward and reverse (For primer sequences see Table S10) and cloned into pET SUMO vector generating a fusion protein with a His-SUMO tag using golden gate cloning (modified from Thermo Fischer). The protein was expressed in BL21-DE3 Rosetta cells at 37 °C. Cells were fractionated (50mM TRIS-HCl pH 7.8, 500mM NaCl, 10mM Imidazole) using Fluidizer (mechanical lysis by high pressure-80 psi). The protein was purified via $Ni^{2+}$ chelate affinity chromatography using a gradient with 200mM imidazole in lysis buffer. Dialysis (50mM Tris-HCl pH 7.8, 500mM NaCl) and SUMO protease digestion for cleavage of the His-SUMO tag were performed simultaneously overnight. The His-SUMO tag was removed from the Tc-FoxQ2 via Re-$Ni^{2+}$ chelate affinity chromatography. Gel-filtration chromatography (Superdex G-30 Healthcare) was performed to remove the remaining contaminations and finally the purified Tc-FoxQ2 was stored in phosphate-buffered saline (PBS). All the steps for purification were done at 4°C. Antibodies were produced in guinea pigs by Eurogentec (Liège, Belgium). The final serum was used straight as the Tc-FoxQ2 antibody with the dilution of 1:1000. Before antibody staining, pre-absorption of anti-FoxQ2 was performed for eliminating non-specific binding.

#### Generation of imaging lines and stocks

*foxQ2-5'-line*

The guide RNAs (gRNAs) were designed with the aid of the flyCRISPR Optimal Target Finder (http://tools.flycrispr.molbio.wisc.edu/targetFinder/; Gratz et al. 2014). The TriGenes gRNA oligo design tool was used for generating the sequences of the oligos to order. The annealed oligos were cloned into the gRNA expression vector p(TcU6b-*BsaI*) via the *BsaI* restriction sites. The detailed annealing and ligation are following the protocol described previously (Gilles et al., 2015). [3xP3:Tc'v-SV40-Cre-2A-EGFP:bhsp68-eb] was designed as a repair template for NHEJ-mediate knock-in by CRISPR/Cas9. For linearizing the plasmid, *Dm-ebony* target site (gRNA-eb) was cloned into this construct (Addgene plasmid # 124068). Each fragment of the construct was amplified by PCR from plasmids available in the laboratory's plasmid library (Table S3) by using primers with overhangs that are complements of two adjacent fragments (Table S10). The 2A-peptide and the target sequences for cleavage were completely added by primers. Overlap extension PCR was performed to assemble all fragments together. In addition, an *ApaI* and a *XbaI* restriction sites were added in end primers for the following ligation. The entire construct was finally cloned into pJET1.2 vector. The helpler plasmid p(bhsp68-Cas9) expressing Cas9 was a gift from Michalis Averof (Addgene plasmid # 65959).

Embryonic injection was performed in *Tc-vermillion$^{white}$* (*Tc-vw*) according to standard procedure (Berghammer et al. 1999; Schinko et al., 2012). Two gRNAs (gRNA1 and gRNA2) targeting the upstream region of *Tc-foxQ2* together with the repair plasmid, Cas9 expression plasmid and gRNA-eb were injected. The final concentration of Cas9 plasmid and the repair plasmid is 500 ng/µl each, and gRNA is 125 ng/µl each. The injected animals were separated into male and female during pupal stage. Each animal was crossed to three *Tc-vw* wild type beetles of the opposite sex. The G1 offspring were screened for black eyes. The transgenic beetle was outcrossed with *Tc-vw* wild type and kept as a new stock.

*rx-5′-up line*

The regulatory region including the endogenous promoter sequence of the gene *Tc-rx* was amplified from genomic DNA by PCR using primer pairs with restriction sites *BamHI* and *NheI* (for primer sequences see Table S10) and cloned into the Dual Promoter pCR®II vector by using the TA Cloning® Kit (Invitrogen). The final construct was designed with the following sections from 5´ to 3´: (1) regulatory regions, (2) endogenous promoter, (3) reporter gene. The reporter gene DsRedExpress was amplified from the other construct by using according primers (Table S10). The construct was designed and created in the vector pslfa1180fa. Afterwards, the cassette [rx-5'up:DsRedEx-SV40] including the regulatory region, promoter, reporter gene, and SV40 was transferred into the piggyBac[3xP3:Tc'v-SV40]fa transformation vector by using the restriction enzymes *AscI* and *FseI* (New England BioLabs). Further steps and treatments for embryonic transgenesis were performed as described (Berghammer et al. 1999; Schinko et al., 2012).
